# Loss of Rer1 causes proteotoxic stress that drives cell competition and inhibits Myc-driven overgrowth

**DOI:** 10.1101/2022.08.17.504145

**Authors:** Pranab Kumar Paul, Rishana Farin S, Wim Annaert, Varun Chaudhary

## Abstract

Cell competition is a developmental phenomenon that allows the selection of healthier cells in a developing tissue. In this process, cells with reduced fitness, conceivably due to harmful mutations, acquire the ‘loser’ status and are eliminated by the fitter (winner) neighboring cells via juxtacrine cell-cell interactions. How various mutations trigger cell competition is an extensively studied question. However, the mechanism of cell competition remains largely elusive. In this study, we reveal previously unknown functions of an ER and Golgi localized protein Rer1 in the regulation of cell competition in the developing *Drosophila* wing epithelium. Our data show that loss of Rer1 leads to the proteotoxic stress marked by the increased phosphorylation of eIF2α. The increased proteotoxic stress in the *rer1* mutant cells led to their elimination via cell competition. Interestingly, we find that Rer1 levels are upregulated upon Myc overexpression, which generates super-competitive cells that overgrow at the expense of the normal neighboring cells. Loss of Rer1 also restricts the growth of Myc-induced super-competitive cells. Moreover, consistent with its known function as a negative regulator of the Notch pathway, our results show that the increased levels of Rer1 in Myc-overexpression led to the downregulation of Notch activity. In summary, these observations provide the first characterization of Rer1 in *Drosophila* and reveal its role in triggering cell competition via the regulation of proteotoxic stress.

## INTRODUCTION

The development of healthy tissue requires the removal of viable but suboptimal cells. In several growing tissues, this is achieved by a specific cell-cell interaction mechanism called cell competition, whereby unfit cells (also called “loser” cells) are eliminated by the surrounding fitter or so-called “winner” cells (Morata and Ripoll, 1975; Simpson, 1979). The best-known example of cell competition is described in the developing *Drosophila* epithelium. Studies have shown that heterozygous mutations in a Ribosomal protein (Rp) gene (also known as *Minute*) give viable flies; however, in a mosaic condition in growing tissue, *Rp*^*+/-*^ cells are eliminated from the tissue when surrounded by the neighboring wild-type (*Rp*^*+/+*^) cells (Abrams, 2002; Milán et al., 2002). While the *Rp*^*+/-*^ mutation affects cellular physiology autonomously, caspase-dependent apoptosis is observed mostly at the boundary between *Rp*^*+/-*^ cells and nearby *Rp*^*+/+*^ cells, which is a hallmark of cell competition (Li and Baker, 2007; Moreno et al., 2002; Oertel et al., 2006). Previous studies have suggested that the loser fate of *Rp*^*+/-*^ cells is due to a reduction in protein translation (Lee et al., 2018; Moreno et al., 2002). However, recently it was shown that the loser fate of Rp^+/-^ is a result of increased proteotoxic stress due to protein aggregation (Albert et al., 2019; Baumgartner et al., 2021; Recasens-Alvarez et al., 2021; Tye et al., 2019).

Moreover, the loser fate is associated with a number of other physiological changes affecting cell fitness, including, 1) reduced metabolic activity due to changes in the levels of mTOR pathway activity (Bowling et al., 2018; Sanaki et al., 2020), 2) loss of apico-basal polarity as a consequence of mutations of the *scribble, dlg*, and *lgl* genes (Brumby and Richardson, 2003), 3) defects in intracellular protein trafficking caused by mutations in the *rab5* gene (Ballesteros-Arias et al., 2014), and 4) deregulation of signaling pathways such as Wnt, BMP, and Hippo (Nagata et al., 2022; Neto-Silva et al., 2010; Vincent et al., 2011). Cells with these perturbations are eliminated through cell competition involving JNK-dependent activation of the proapoptotic factors (Moreno et al., 2002; Pinal et al., 2019). Furthermore, several perturbations can also provide a competitive advantage to the cells over their wild-type neighbors. For example, the overexpression of a proto-oncogene *Myc*, a master regulator of cell proliferation and growth, increases the relative fitness of the cells. Thus, clonal expression of Myc in a developing tissue generates super-competitor cells, which proliferate at the expense of the wild-type neighbors (de la Cova et al., 2004; Moreno and Basler, 2004). Myc protein plays a broad role in promoting cell growth and proliferation, which ranges from the regulation of gene expression to protein synthesis and metabolism (de la Cova et al., 2014; Eisenman, 2001; Johnston et al., 1999). Studies have shown that Myc overexpression leads to proteotoxic stress due to increased protein synthesis (Iritani and Eisenman, 1999; Nguyen et al., 2018; Tameire et al., 2015). Furthermore, Myc-driven overgrowth is dependent on the activation of the cytoprotective unfolded protein response pathways (UPR) (Nagy et al., 2013). This includes PERK-mediated phosphorylation of eukaryotic initiation factor 2 alpha (eIF2α) and induction of autophagy to reduce protein translation and clear misfolded proteins, respectively (Levy et al., 2017; Zhang et al., 2020). However, a clear understanding of how Myc and UPR work together to promote a proliferative environment within a cell is lacking.

Here, we have studied the role of Retention in Endoplasmic Reticulum-1 (Rer1) protein in the competitive cell proliferation in the developing *Drosophila* wing epithelium. Mutations in the *rer1* gene were first described in yeast, where it was identified in a screen as a factor required for proper transport of Sec12p between ER and Golgi (Nishikawa and Nakano, 1993). Later studies showed that Rer1 protein localizes to ER and cis-Golgi compartments (Sato et al., 1995) and that it is required for the COPI-dependent retrieval of various ER-resident proteins, including Sec12p, Sec63p and Sec71p from the Golgi (Füllekrug et al., 1997; Sato et al., 1997, 2001). Furthermore, studies have shown that Rer1 is also required for the assembly of multisubunit protein complexes, for example, the γ-secretase complex, yeast iron transporter and skeletal muscle nicotinic acetylcholine receptor (nAChR) (Kaether et al., 2007; Park et al., 2012; Sato et al., 2004; Spasic et al., 2007; Valkova et al., 2011). The γ-secretase complex consists of four subunits, nicastrin (NCT), presenilin (PS), anterior pharynx defective 1 (APH-1) and presenilin enhancer 2 (PEN-2) and it is involved in the processing of several transmembrane proteins (type-I) such as amyloid-β (Aβ) peptide APP and the Notch receptor (De Strooper, 2003; Jurisch-Yaksi et al., 2013). Rer1 has been shown to interact with the subunits NCT and PEN2, and regulate the assembly of the γ-secretase complex (Kaether et al., 2007; Spasic et al., 2007). However, the role of Rer1 in Notch signaling activity is unclear and appears to be context dependent. On the one hand, Rer1 was shown to prevent γ-secretase assembly and therefore act as a negative regulator of the Notch pathway (Jurisch-Yaksi et al., 2013; Kim et al., 2018). While on the other, it was shown that loss of Rer1 led to the degradation of the γ-secretase subunits with a concomitant loss of Notch-dependent maintenance of the neural stem cells (Hara et al., 2018), thus acting as a positive regulator. Despite the fact that Rer1 is evolutionarily conserved from yeast to mammals (Annaert and Kaether, 2020; Füllekrug et al., 1997; Ghavidel et al., 2015), its function in the development of organisms remains largely unknown (Annaert and Kaether, 2020).

In this study, we have characterized the role of Rer1 in the development of *Drosophila* wing epithelium. By creating a *rer1* loss-of-function mutant, we show that *rer1* is an essential gene in *Drosophila* and loss of Rer1 causes cell death. More importantly, we find that *rer1* mutant cells attain the “loser” fate and are eliminated specifically via the process of cell competition. We further show that the loser fate of *rer1* mutant cells is due to an increase in proteotoxic stress. We have also analyzed the role of Rer1 in Myc-induced super-competition and Rer1 levels were found to be upregulated upon Myc overexpression, which further led to downregulation of Notch activity. More importantly, we find that loss of Rer1 function is sufficient to reduce Myc-induced overgrowth. In summary, our results demonstrate that Rer1 is an essential protein for proper maintenance of protein homeostasis and competitive cell survival in a developing tissue.

## RESULTS

### *rer1* is an essential gene for the development

We first set out to characterize the role of Rer1 during *Drosophila* development. To this end, we generated a *rer1* knockout mutant by imprecise excision of a p-element insertion in the *rer1* locus (see materials and methods). A loss-of-function mutation in *rer1* containing a 1560bp deletion in the coding region was identified (**Supplemental Figure 1A**). Quantitative RT-PCR analysis in the homozygous mutant animals showed a complete loss of *rer1* mRNA levels, indicating that the deletion in *rer1* caused a complete loss-of-function (**Supplemental Figure 1B**). Further analysis showed that the homozygous *rer1* mutants (*rer1*^*–/–*^) failed to develop into pupa and died during larval stages (**Supplemental Figure 1C**). However, to rule out that the lethality could be due to a second site mutation in another essential gene, we performed rescue experiments by using a genomic-rescue construct expressing GFP-tagged Rer1 via the endogenous promoter. The expression of GFP-Rer1 in homozygous *rer1*^*–/–*^ led to a complete rescue of lethality, confirming the specificity of the mutant (**Supplemental Figure 1C**). These results demonstrate that *rer1* is an essential gene in *Drosophila* development which encouraged us to further analyze the importance of Rer1 at the tissue level.

To this end, we first depleted Rer1 in the posterior compartment of the developing *Drosophila* wing imaginal discs by expressing *rer1*-RNAi using the *hedgehog (hh)-Gal4* driver (**Supplemental Figure 1D**). To test the efficiency of the knockdown, we expressed *rer1*-RNAi in the *GFP-rer1* genomic-rescue flies. Here we observed a strong downregulation of the GFP-Rer1 levels (**Supplemental Figure 1E**), suggesting that *rer1*-RNAi effectively downregulated the Rer1 levels. We next tested the effect of Rer1 depletion on cell death by analyzing the expression of cleaved Death caspase-1 (Dcp-1) and Acridine Orange (AO), which act as markers for apoptosis (see materials and methods). The depletion of Rer1 in the posterior compartment led to a strong increase in both Dcp-1 and AO positive cells compared to the control anterior compartment (**Supplemental Figure 1F-J**).

To further confirm these results, we analyzed the Dcp-1 levels in *rer1*^*–/–*^ mutant clones, generated using the Flippase (FLP)-Flp recognition target (FRT)-system (see materials and methods). *rer1*^*–/–*^ mutant clones were induced using heat-shock Flippase during early larval stages, and 96 hrs after heat-shock (AHS) third-instar wing discs were stained with the anti-cleaved Dcp-1 antibody. Consistent with the RNAi experiment, *rer1*^*–/–*^ mutant clones showed upregulation of Dcp-1 levels (**Figure 1A and B**). Furthermore, the *rer1*^*–/–*^ mutant clones did not grow to the same extent as the control *rer1*^*+/+*^ (wild-type) clones generated at the same time (**Figure 1, compare RFP negative area in A and B**), and both Dcp-1 levels and clone size were rescued by the expression of GFP-Rer1 (**Figure 1D**). Interestingly, we also observed that Dcp-1 positive cells were present specifically at the boundary of *rer1*^*–/–*^ cells and *rer1*^*+/+*^ cells (**Figure 1C, blue-arrows**). Previous studies have shown similar boundary-restricted cell death which is specific to cell competition (Kale et al., 2015; Lolo et al., 2012; Nagata et al., 2019; Sanaki et al., 2020). Based on this, we hypothesized that loss of Rer1 triggers cell competition, whereby cells lacking Rer1 are identified as unfit or loser cells and are eliminated by the adjacent wild-type (winner) cells.

**Figure 1:**
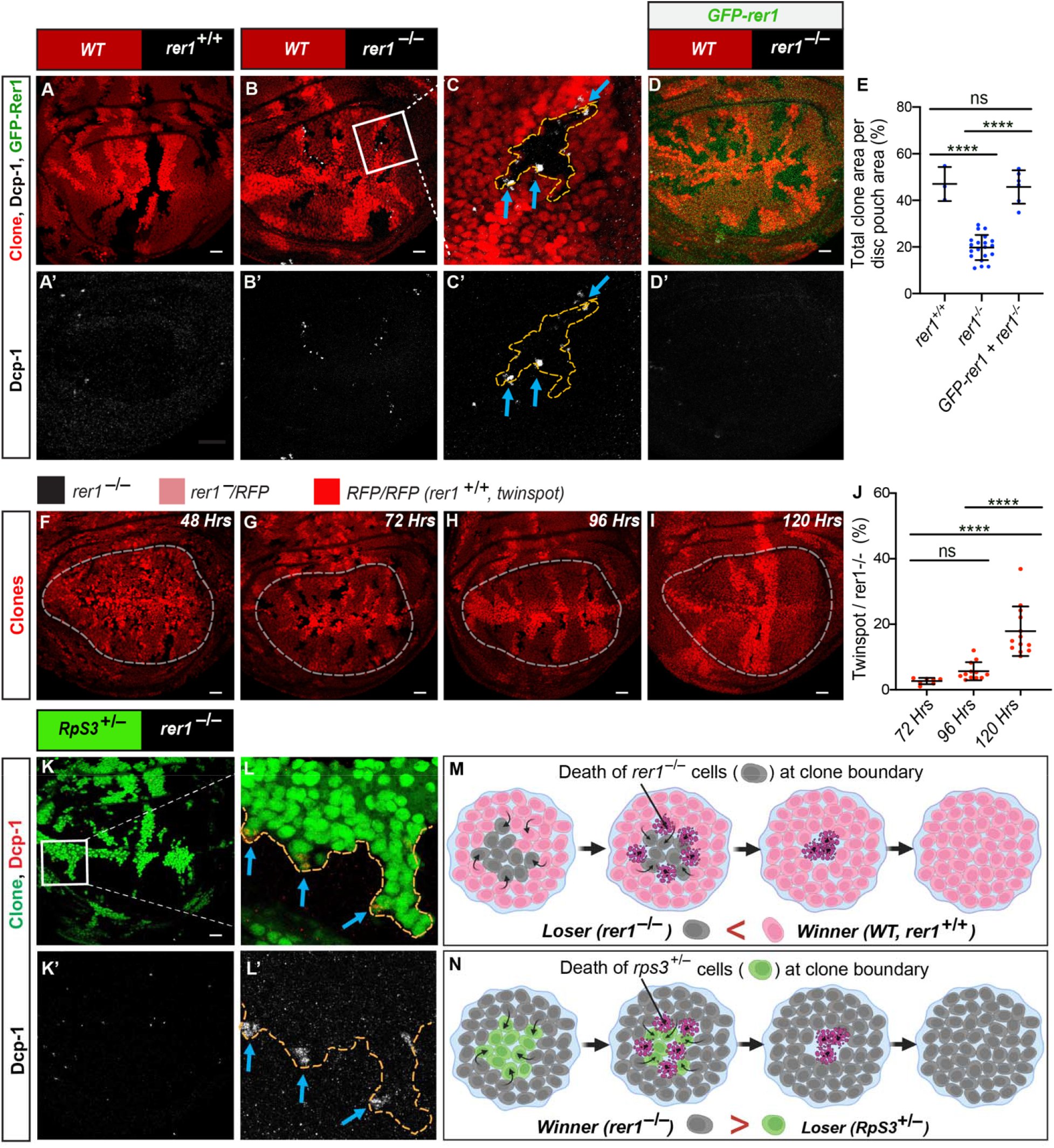
*rer1^−/−^* cells are eliminated through cell competition. (A-C) Third-instar larval wing epithelium with hs-FLP-induced mitotic clones of (A) WT (wild-type; rer1^+/+^), and (B) rer1^−/−^ genotypes, immuno-stained for the anti-cleaved Dcp-1. (C) A magnified image of the inset (white box) in B. (D) rer1^−/−^ mutant clones in GFP-rer1 background stained with anti-cleaved Dcp-1. (E) Quantification of the RFP-negative clones’ area in WT (rer1^+/+^; N=3); mutant (rer1^−/−^, N=20) and rescue (rer1^−/−^+ GFP-rer1, N=6). Statistical analysis was performed using the Ordinary one-way ANOVA with Tukey’s multiple comparison test. (F-I) Wing imaginal disc harboring rer1^−/−^ mutant clones induced by hs-FLP at 48, 72, 96 and 120 hrs prior to dissection of third-instar larvae. RFP-negative (black) represents rer1^−/−^, lighter red marks heterozygous rer1^−^/RFP, and brighter red areas represent RFP/RFP (rer1^+/+^; twin-spot). (J) The relative size of twin-spots (RFP/RFP) versus mutant (RFP negative) areas at 72 hrs (N=7), 96 hrs (N=12), and 120 hrs (N=14), measured within the white dotted lines. Twin spots at 48 hrs were not quantified because of higher patchiness. Statistical analysis was performed using the Ordinary one-way ANOVA with Tukey’s multiple comparison test. (K) hs-FLP-induced heterozygous RpS3^+/–^ mutant clones (GFP-positive cells) juxtaposed to rer1^−/−^ cells (GFP negative), and immuno-stained for the anti-cleaved Dcp-1. (L) A magnified image of the inset in K (white box). (M-N) Schematic diagram illustrating the concept of winner and loser fate in cell competition between WT (rer1^+/+^) and rer1^−/−^ tissues; and rer1^−/−^ and RpS3^+/–^ tissues, respectively. The interfaces between the clone areas are marked with yellow dotted lines. SB= 20 μm.

Previous studies have shown that the elimination of loser cells requires activation of the JNK pathway (Moreno et al., 2002). Thus, we next asked if the loser fate and the elimination of *rer1*^*–/–*^ cells are JNK dependent. We first checked if the JNK pathway was active in the *rer1*^*–/–*^ cells, by analyzing the expression of a JNK target gene, *head involution defective* (*hid*), using the *hid-LacZ* reporter. As shown in **Supplemental Figure 2A-C**, *hid-LacZ* levels were upregulated in *rer1*^*–/–*^ cells, indicating activation of JNK signaling. Next, we asked if the activation of JNK activity contributed to the elimination of the *rer1*^*–/–*^ mutant cell. To this end, we expressed dominant-negative Basket (Bsk), the ortholog of JNK in *Drosophila* (Riesgo-Escovar et al., 1996), in *rer1*^*–/–*^ mutant cells. Analysis of the clone area showed that the expression of dominant-negative Basket modestly rescued *rer1*^*–/–*^ mutant cells from elimination (**Supplemental Figure 2D-G**), suggesting that JNK activity is required for the elimination of *rer1*^*–/–*^ mutant cells.

### *rer1*^*–/–*^ cells are eliminated through cell competition

We further tested if the elimination of *rer1*^*–/–*^ cells occurs via cell competition by analyzing two other hallmark features; (1) the growth rate difference of the loser cells in mosaic analysis; and (2) the changing of the fate of mutant cells as loser/winner depending upon the relative fitness with the neighboring cells (Colom et al., 2020; Ellis et al., 2019). To verify the extent of differential growth rate, we compared the areas occupied by *rer1*^*+/+*^ (RFP/RFP, also called twin spot) and *rer1*^*–/–*^ (RFP negative) clones in the developing wing imaginal disc at various post-induction time points. Consistent with the increased cell death observed in *rer1*^*–/–*^ mutant clones, we find that the proportion of twin spots increased significantly over time compared to the *rer1*^*–/–*^ area (**Figure 1F-I, quantified in J**).

Next, we tested if the fate of *rer1*^*–/–*^ cells could be reversed due to the relative fitness of the neighboring cells. To this end, we generated *rer1*^*–/–*^ clones along with ribosomal protein S3 (*RpS3)* mutant clones in the same imaginal disc, causing juxtaposition of *rer1*^*–/–*^ cells with the *RpS3*^*+/–*^ cells instead of the normal wild-type cells. Homozygous *RpS3* mutant cells fail to survive on their own, while heterozygous *RpS3* mutant (*RpS3*^*+/–*^) cells are eliminated by surrounding wild-type cells (*RpS3*^*+/+*^) via cell competition (Moreno et al., 2002). Here, we observe a dramatic change in the growth of *rer1*^*–/–*^ mutant clones, which were significantly larger than the *RpS3*^*+/–*^ mutant clones, and the Dcp-1 staining could now be observed in the *RpS3*^*+/–*^ mutant cells at the boundary (**Figure 1K-L, illustrated in M and N**). This suggests that activation of cell death in *rer1*^*–/–*^ could be reversed by reducing the fitness level of the neighbors. Altogether, these results confirm that the removal of the *rer1*^*–/–*^ cells occurs via competitive cell-cell interactions.

### Deficiency of Rer1 creates proteotoxic stress

Next, we asked what physiological changes are manifested in the cells upon the loss of Rer1, which led to its loser fate. Previous studies have shown that Rer1 is localized dynamically between the ER and cis-Golgi compartments in *Saccharomyces cerevisiae* (Sato et al., 1995, 2001), and mammalian cells (Füllekrug et al., 1997; Spasic et al., 2007). However, its localization in *Drosophila* remains unknown. To ascertain if *Drosophila* Rer1 has a similar function, we first checked its localization using the GFP-Rer1 genomic rescue construct. Consistent with its known localization, we find that the GFP-Rer1 was colocalized with both ER and Golgi markers in the wing epithelial cells (**Supplemental Figure 3A-B**). As Rer1 is known for its function in the protein quality control process, we next asked if the loss of Rer1 also affected protein homeostasis in flies. Thus, we analyzed the level of phosphorylated eIF2α (p-eIF2α), which is a well-known marker of proteotoxic stress (Pakos-Zebrucka et al., 2016). *rer1*^*–/–*^ mutant clones showed upregulation of p-eIF2α levels compared to the control region (**Figure 2A and B**), indicating activation of the UPR pathway. The increased level of p-eIF2α in *rer1*^*–/–*^ mutant clones was rescued upon the expression of GFP-Rer1 (**Supplemental Figure 4A-E; quantified in C and F**).

**Figure 2:**
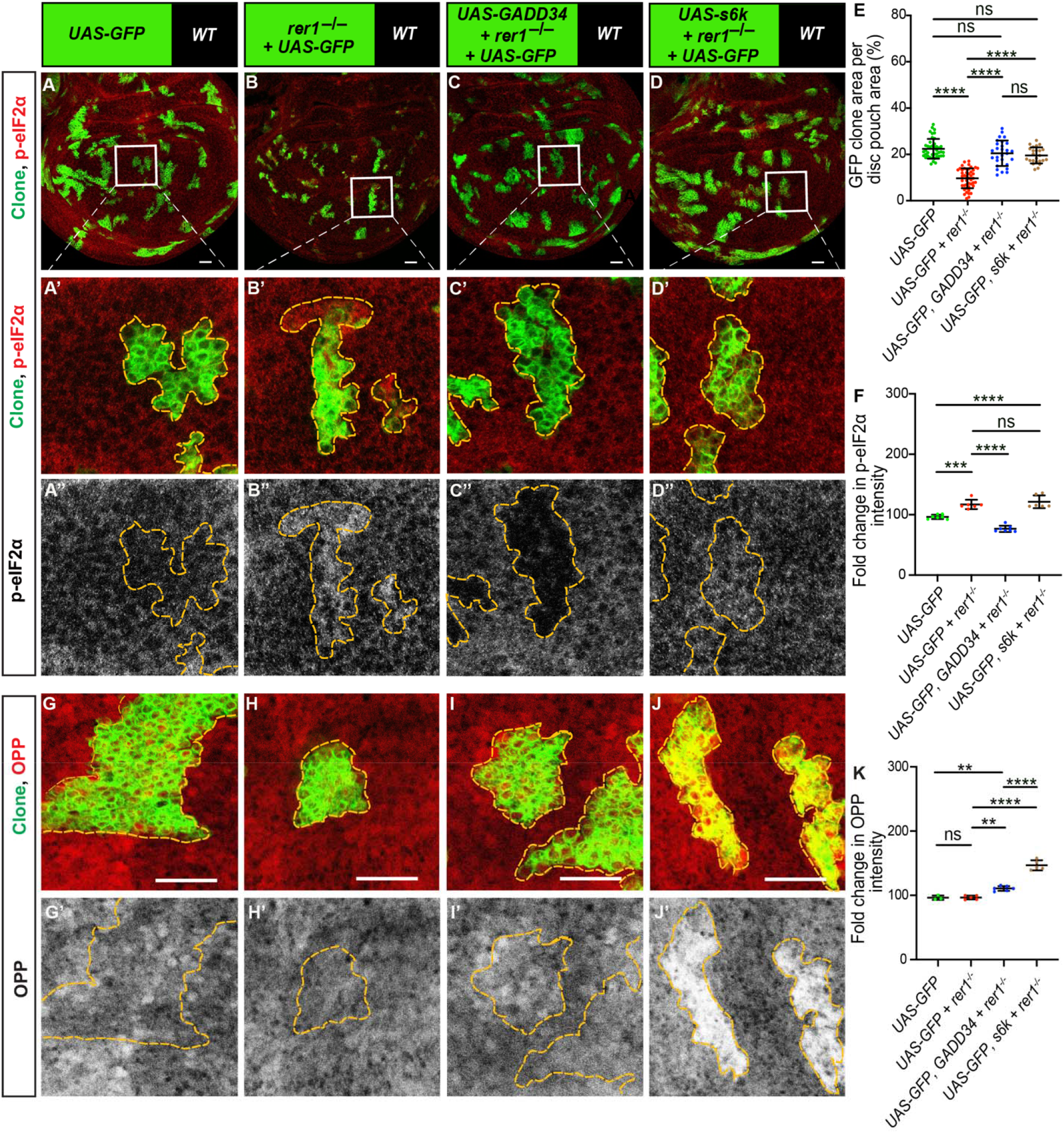
Loss of *rer1* function creates proteotoxic stress on the wing disc epithelium. (A-C) Third-instar wing epithelium containing hs-FLP-induced MARCM clones of (A) UAS-GFP (rer1^+/+^), (B) UAS-GFP, rer1^−/−^, (C) UAS-GFP, rer1^−/−^ + UAS-GADD34 and, (D) UAS-GFP, rer1^−/−^ + UAS-s6k genotypes, stained with the anti-p-eIF2α antibody. (A’-D’) Magnified images of the insets in A, B, C and D are shown in A’, B’, C’ and D’, respectively. (E) Quantification of the relative size of GFP-labeled clones area in UAS-GFP (rer1^+/+^) (A, N=42), UAS-GFP, rer1^−/−^, (B, N=46), UAS-GFP, rer1^−/−^ + UAS-GADD34 (C, N=28) and, UAS-GFP, rer1^−/−^ + UAS-s6k (D, N=24). (F) Quantification of the p-eIF2α levels inside the GFP positive clones with respect to the nearby GFP negative control tissue in, UAS-GFP (rer1^+/+^) (A, N=7), UAS-GFP, rer1^−/−^ (B, N=6), UAS-GFP, rer1^−/−^ + UAS-GADD34 (C, N=7) and, UAS-GFP, rer1^−/−^ + UAS-s6k (D, N=6). (G-J) OPP assay on third-instar discs containing hs-FLP-induced MARCM clones of (G) UAS-GFP (rer1^+/+^), and (H) UAS-GFP, rer1^−/−^, (I) UAS-GFP, rer1^−/−^ + UAS-GADD34 and, (J) UAS-GFP, rer1^−/−^ + UAS-s6k genotypes. (K) Quantification of the signal intensity of OPP inside the GFP positive clones with respect to nearby GFP negative control tissue in UAS-GFP (G, N=3), UAS-GFP, rer1^−/−^ (H, N=5), UAS-GFP, rer1^−/−^ + UAS-GADD34 (I, N=6) and, UAS-GFP, rer1^−/−^ + UAS-s6k (J, N=5). Borders between the GFP positive and GFP negative areas are marked with yellow dotted lines. Statistical analyses in E, F and K were performed using the Ordinary one-way ANOVA with Tukey’s multiple comparison test. SB = 20 μm.

Being an essential component of the translational machinery, phosphorylation of eIF2α has been associated with a reduction in protein synthesis (Pakos-Zebrucka et al., 2016). We next tested whether higher levels of p-eIF2α in *rer1*^*–/–*^ mutants also affected global protein translation, which perhaps led to their elimination. To test this, we performed an O-propargyl-puromycin (OPP) incorporation assay, where OPP, an analog of puromycin, is incorporated into nascent polypeptide chains directly allowing rapid assessment of global protein synthesis (Schmidt et al., 2009), (see materials and methods). Surprisingly, in these experiments, *rer1*^*–/–*^ cells did not show significant changes in OPP levels (**Figure 2G and H**). Although we can not rule out that there were mild effects that were undetectable with the OPP assay, these results indicated that the global translation was not strongly affected by the loss of Rer1.

Irrespective, we next questioned whether phosphorylation of eIF2α is the reason for the loser fate of the *rer1* mutant cells leading to their elimination. To test this, we overexpressed growth arrest and DNA damage-inducible 34 protein (GADD34), which provides specificity to protein phosphatase 1 for the dephosphorylation of p-eIF2α (Novoa et al., 2001). As expected, the overexpression of GADD34 in *rer1*^*–/–*^ mutant cells led to a strong reduction in the p-eIF2α levels (**Figure 2C, C’’**) and an increase in the OPP incorporation (**Figure 2I, I’**). More importantly, the expression of GADD34 rescued the growth of *rer1*^*–/–*^ mutant clones (**Figure 2E**), suggesting that the competitive elimination of the *rer1*^*–/–*^ cells is due to the phosphorylation of eIF2α.

Next, we asked whether increasing protein translation independently of p-eIF2α dephosphorylation would affect the growth of *rer1*^*–/–*^ mutant cells. We expressed S6K, which is a target of the mTOR pathway and has been shown to cause an increase in protein synthesis (Hay and Sonenberg, 2004; Jefferies et al., 1997; Neufeld et al., 1998; Shima et al., 1998). Expression of S6K in *rer1*^*–/–*^ cells did not cause a significant change in the levels of p-eIF2α (**Figure 2D, D’’; quantified in F**), indicating the proteotoxic stress in *rer1*^*–/–*^ cells was not affected by the expression of S6K. However, OPP levels were higher and the growth of *rer1*^*–/–*^ cells was also rescued by the expression of S6K (**Figure 2J, J’; quantified in K and E**). These results suggest that increasing protein translation could also provide a competitive advantage to the *rer1* mutant cells.

### Rer1 is required for Myc-induced super-competition

We next asked whether Rer1 would be involved in other cell competition pathways which have been associated with the activation of UPR. Studies have shown that overexpression of Myc provides a growth advantage to the cells, which attain the super-competitor status that leads to the proliferation of the Myc-overexpressing cells at the expense of the neighboring wild-type cells (de la Cova et al., 2004; Moreno and Basler, 2004). Furthermore, UPR activation is required for Myc-driven overgrowth (Nagy et al., 2013). Thus, we tested if Rer1 plays a role in the growth of Myc-overexpressing cells. To this end, we generated Myc overexpressing super-competitive clones in the *Drosophila* wing imaginal disc, in either wild-type (**Figure 3B**) or *rer1*^*–/–*^ mutant background (**Figure 3C**). As shown in Figure 3, the overgrowth phenotype observed due to the overexpression of Myc was reduced in the *rer1*^*–/–*^ cells (**Figure 3A-D, quantified in E; compare B and C**). These results suggest that Rer1 is required for Myc-induced overproliferation.

**Figure 3:**
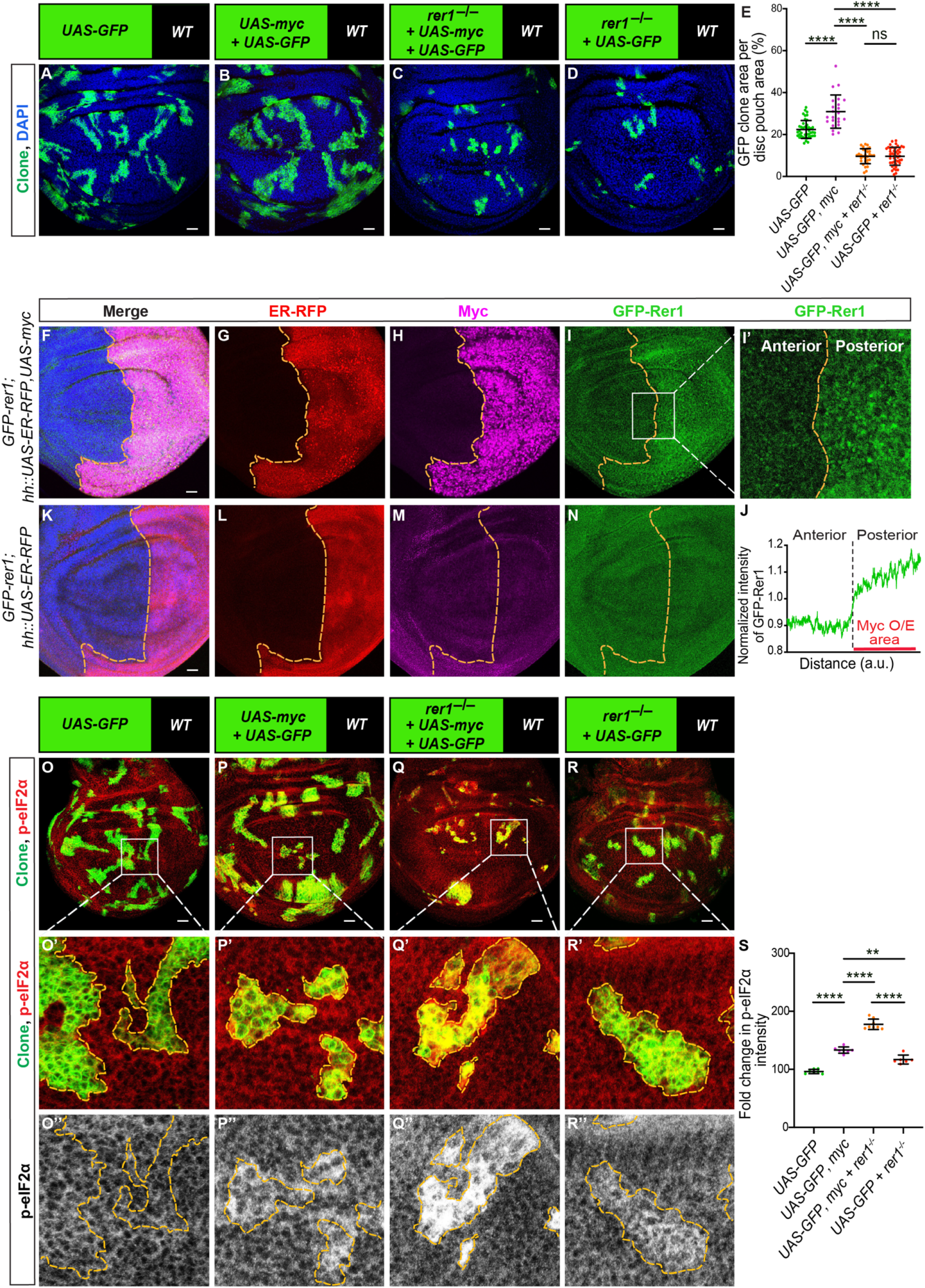
Rer1 is required for Myc-induced super-competition. (A-D) Third-instar discs wing epithelium discs containing hs-FLP-induced MARCM clones of (A) UAS-GFP (rer1^+/+^), (B) UAS-GFP, UAS-myc, (C) UAS-GFP, rer1^−/−^ + UAS-myc and, (D) UAS-GFP, rer1^−/−^ genotypes, stained with DAPI to label the nuclei. (E) Quantification of the relative size of GFP-labeled clones area in UAS-GFP (rer1^+/+^) (A, N=42), UAS-GFP, UAS-myc, (B, N=23), UAS-GFP, rer1^−/−^ + UAS-myc (C, N=31) and, UAS-GFP, rer1^−/−^ (D, N=46). (F-N) GFP-rer1 levels were analyzed in the posterior compartment upon hh-Gal4 mediated Myc overexpression [GFP-rer1 + hh::UAS-ER-RFP + UAS-myc, (F-I)] and compared to the anterior compartment relative to the control wing imaginal disc [GFP-rer1 + hh::UAS-ER-RFP, (K-N)]. (I’) A magnified image of the inset (white box) in I. (J) Quantification of the level of GFP tagged Rer1 in I’. ‘N’ for experimental and control discs are 10 and 9, respectively. (O-R) Third-instar discs containing hs-FLP-induced MARCM clones of (O) UAS-GFP (rer1^+/+^), (P) UAS-GFP, UAS-myc, (Q) UAS-GFP, rer1^−/−^ + UAS-myc and, (R) UAS-GFP, rer1^−/−^ genotypes, stained with the anti-p-eIF2α antibody. (O’-R’) Magnified image of the insets (white box) in O, P, Q and R are shown in O’, P’, Q’ and R’, respectively. (S) Quantification of the p-eIF2α levels inside the GFP positive clones with respect to the nearby GFP negative control tissue in, UAS-GFP (rer1^+/+^) (O, N=7), UAS-GFP, UAS-myc, (P, N=7), UAS-GFP, rer1^−/−^ + UAS-myc (Q, N=8) and, UAS-GFP, rer1^−/−^ (R, N=6). Borders between the GFP positive and GFP negative areas are marked with yellow dotted lines. Statistical analyses in E and S were performed using the Ordinary one-way ANOVA with Tukey’s multiple comparison test. SB=20 μm.

Since Myc overexpression causes increased levels of gene expression and protein translation (Iritani and Eisenman, 1999), we asked whether Rer1 levels are modulated to maintain protein homeostasis. To test this, we utilized the GFP-Rer1 genomic-rescue line to detect changes in Rer1 expression. We observed higher levels of GFP-Rer1 in discs overexpressing Myc in the posterior compartment via the *hh-GAL4* driver (**Figure 3F-I’, quantified in J**), indicating that higher levels of Rer1 might be required for the maintenance of proper protein homeostasis upon Myc overexpression. We further strengthened these conclusions, by analyzing the UPR upon Myc overexpression in the presence and absence of Rer1. As expected, we observed higher levels of p-eIF2α in *rer1*^*–/–*^ mutant cells overexpressing Myc (**Figure 3Q-Q’’**) as compared to either *rer1*^*–/–*^ mutants (**Figure 3R-R’’**) or Myc overexpression alone (**Figure 3P-P’’**). Altogether, these results show that higher levels of Rer1 in Myc-overexpression partially alleviated protein stress supporting Myc-driven overgrowth.

### Rer1 acts as a negative regulator of the Notch pathway

We next asked if the increased levels of Rer1 upon Myc-overexpression affected specific protein or signaling pathways. Thus, we turned our attention to the known targets of Rer1. Past studies have established a strong connection between Rer1 and the Notch pathway. As Rer1p negatively regulates γ-secretase assembly (Spasic et al., 2007), morpholino-mediated knockdown of Rer1 in zebrafish larvae augmented Notch signaling (Jurisch-Yaksi et al., 2013). While this was confirmed in the murine embryonic neural progenitor cells (Kim et al., 2018), another study also in the murine neural stem cells and the cerebrum showed a positive effect of Rer1 on the Notch pathway (Hara et al., 2018; Kim et al., 2018; Annaert and Kaether 2020). Nevertheless, whether *Drosophila* Rer1 is involved in Notch pathway regulation is not known. Therefore, we first analyzed Notch signaling activity upon loss of Rer1. We tested the levels of the Notch intracellular domain (NICD), which is released by γ-secretase following ectodomain shedding of the Notch receptor (Jurisch-Yaksi et al., 2013; Strooper et al., 1999) and acts as a transcription activator of the Notch pathway (Borggrefe and Oswald, 2009; Ross and Kadesch, 2001). Here we observe that only a few *rer1*^*–/–*^ mutant clones showed a modest increase in NICD levels (**Supplemental Figure 5A-D, and quantified in E**), indicating that Rer1 could act as a negative regulator of the Notch pathway. However, since we observed only a mild effect with low penetrance, we aimed to further validate the effect of Rer1 on the Notch pathway. Thus, we asked whether the increase in Rer1 levels, as observed upon Myc overexpression, would be sufficient to modulate Notch activity. Interestingly, Myc-expressing clones showed lower levels of NICD as compared to the neighboring wild-type cells (**Figure 4A and B, quantified in E**), and a concomitant decrease in the Notch target gene Cut (**Figure 4G, G’, quantified in G’’**). This suggests that Myc-overexpression caused downregulation of Notch activity. Next, we tested if loss of Rer1 modulated Notch signaling in Myc expressing cells. *rer1*^*– /–*^ mutant clones with Myc-overexpression showed normal NICD levels (**Figure 4C and D, quantified in E**) and a rescue of Cut expression (**Figure 4H and I, quantified in H’’**).

**Figure 4:**
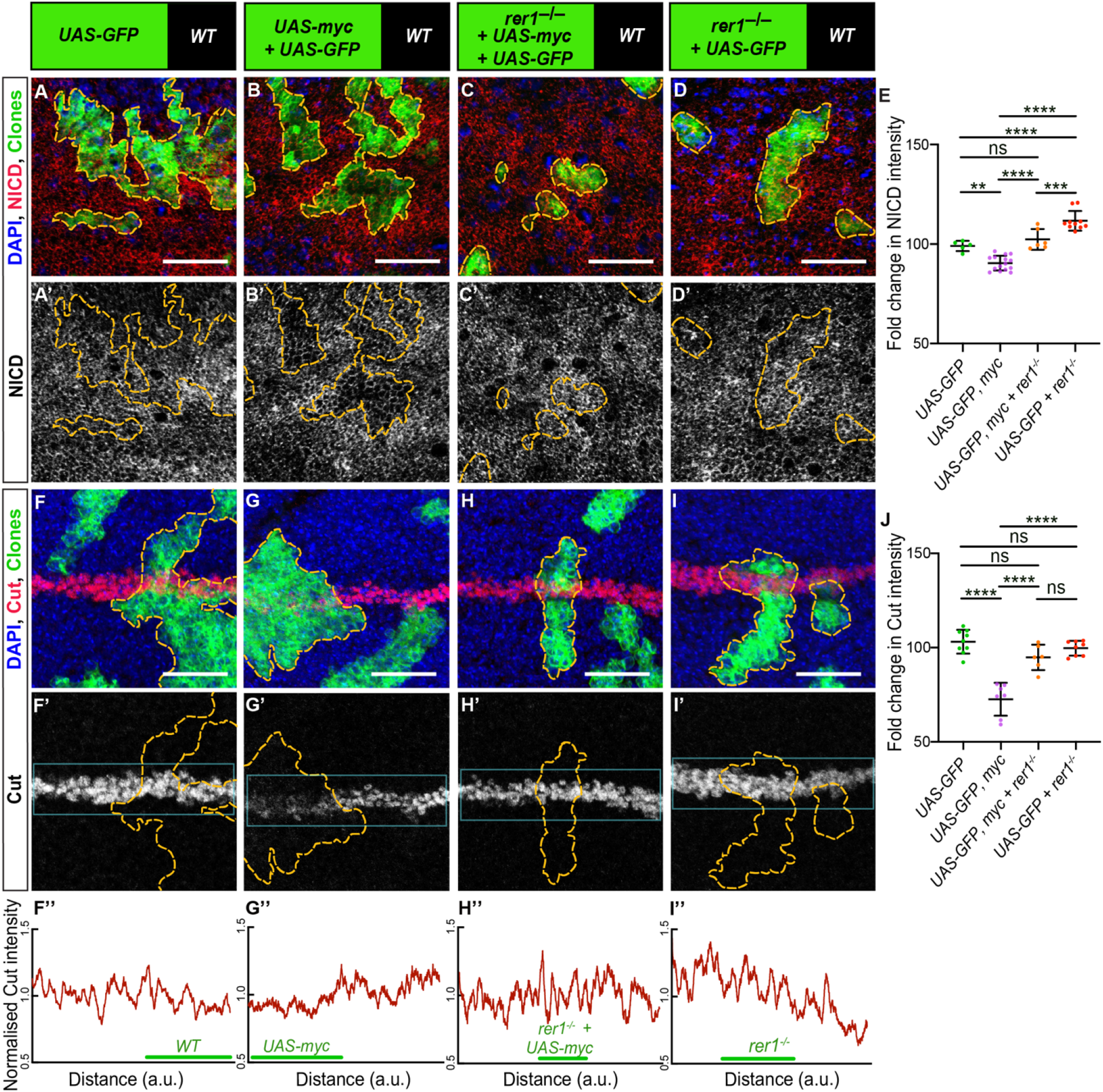
Myc overexpression causes loss of Notch pathway activity. (A-D) Third-instar discs containing hs-FLP-induced MARCM clones of (A) UAS-GFP (rer1^+/+^), (B) UAS-GFP, UAS-myc, (C) UAS-GFP, rer1^−/−^ + UAS-myc and, (D) UAS-GFP, rer1^−/−^ genotypes, immuno-stained for the NICD (red) with nuclei labeled in blue. (E) Quantification of the NICD levels inside the GFP positive clones with respect to the nearby GFP negative control tissue in, UAS-GFP (rer1^+/+^) (A, N=5), UAS-GFP, UAS-myc (B, N=15), UAS-GFP, rer1^−/−^ + UAS-myc (C, N=6) and, UAS-GFP, rer1^−/−^ (D, N=10). (F-I) Third-instar discs containing hs-FLP-induced MARCM clones of (F) UAS-GFP (rer1^+/+^), (G) UAS-GFP, UAS-myc, (H) UAS-GFP, rer1^−/−^ + UAS-myc and, (I) UAS-GFP, rer1^−/−^ genotypes, immuno-stained for the Cut (red) with nuclei labeled in blue. (J) Quantification of the Cut levels inside the GFP positive clones with respect to the nearby GFP negative control tissue in, UAS-GFP (rer1^+/+^) (F, N=9), UAS-GFP, UAS-myc, (G, N=7), UAS-GFP, rer1^−/−^ + UAS-myc (H, N=6) and, UAS-GFP, rer1^−/−^ (I, N=8). (F’’-I’’) Cut intensities inside the insets (Cyan boxes) in F’, G’, H’ and I’ are normalized and quantified in F’’, G’’, H’’ and I’’, respectively. The green lines along the distance (X-axis) of the graph marked the area of GFP positive clones with the respective genotypes in green. Borders between the GFP positive and GFP negative areas are marked with yellow dotted lines. Statistical analyses in E and J were performed using the Ordinary one-way ANOVA with Tukey’s multiple comparisons test. SB=20 μm.

Altogether, these results show that in the *Drosophila* wing epithelium, Rer1 plays a negative role in the regulation of the Notch pathway, and Myc overexpression causes Rer1-dependent downregulation of Notch activity. It will be further interesting to assess whether modulation of Notch activity affected Myc-driven overgrowth.

## DISCUSSION

Our study provides the first characterization of the role of Rer1 during *Drosophila* development, wherein we demonstrate that Rer1 plays an essential role in competitive cell proliferation by maintaining proper cellular protein homeostasis. Loss of Rer1 triggers proteotoxic stress, which leads to reduced fitness of the cells and therefore the loser fate. Our work supports the finding of several recent studies, which have also shown that proteotoxic stress caused by the loss of various factors is the primary cause of the loser fate in competitive cell interactions. For instance, mutations in genes encoding RNA Helicase *Hel25E*, E3 ubiquitin ligase *Mahjong* and various Ribosomal proteins, were shown to have higher levels of proteotoxic stress, causing an increase in the levels of p-eIF2α (Baumgartner et al., 2021; Kiparaki et al., 2022; Langton et al., 2021; Nagata et al., 2019; Ochi et al., 2021; Recasens-Alvarez et al., 2021).

Phosphorylation of eIF2α is normally associated with the reduction in global protein translation (Boye and Grallert, 2020; Hinnebusch and Lorsch, 2012; Wek, 2018). However, paradoxically, we find that increased p-eIF2α in *rer1* mutant clones did not lead to a significant reduction in global protein synthesis, as measured by OPP incorporation. While these results are consistent with the other reports where phosphorylation of eIF2α did not lead to a reduction in the global protein translation (Dever, 2002; Stonyte et al., 2018), it is also conceivable that *rer1* mutants only have a partial effect on the phosphorylation of eIF2α and therefore protein translation, which was undetectable with the OPP assay. Nevertheless, the loser status of the *rer1* mutant cells with almost normal translation is consistent with similar observations for other cell competition factors by recent studies (Baumgartner et al., 2021; Kiparaki et al., 2022). Interestingly, we also find that increasing translation either by the dephosphorylation of eIF2α via GADD34 or by the activation of the TOR pathway improved the fitness of *rer1* mutant cells. It is possible that a specific protein or a set of proteins were affected in the *rer1* mutant cells, which were restored by increasing translation.

An important finding in our study is the increased levels of Rer1 observed upon Myc-overexpression. Previous studies have shown that Myc-overexpression leads to the activation of the UPR pathways, whereby PERK-mediated phosphorylation of eIF2α is observed (Nguyen et al., 2018). The increase in p-eIF2α was shown to play a cytoprotective role which is required for the growth of Myc expression tumors in mice (Nguyen et al., 2018; Zhang et al., 2020). Our results show that Myc-expressing cells failed to grow in the absence of Rer1 even with higher levels of p-eIF2α, indicating that Rer1 is essential for Myc-driven overgrowth. It is tempting to speculate that Rer1 may also be part of UPR pathways that could work independently of the p-eIF2α mediated block in global protein translation. Targeting Myc protein directly as a therapeutic approach has not been successful so far (Chen et al., 2018), therefore, our results provide a strong justification to further analyze the role of Rer1 for potential therapeutic approaches targeting Myc tumors.

Moreover, our results also confirm the previously suggested role of Rer1 as a negative regulator of the Notch pathway. Rer1 has been shown to negatively impact the assembly of the γ-secretase complex (Spasic et al, 2007) and thus its levels in the post-Golgi compartments, where it acts in the ligand-dependent cleavage of Notch receptors leading to the activation of the Notch pathway. We find that loss of Rer1 in an otherwise wild-type background did not show a strong effect on the Notch pathway activity, which was expected to be upregulated. However, wing epithelial cells already have high Notch activity (de Celis et al., 1996; Go et al., 1998), which in turn suggests that high levels of γ-secretase would be present at the membrane. Therefore, it is possible that the effect of loss of Rer1 is limited by the availability of additional complex proteins to be recruited at the membrane.

Consistent with its role as a negative regulator of the Notch pathway (Jurisch-Yaksi et al, 2013; Kim et al., 2018), we find that higher levels of Rer1 observed upon Myc overexpression caused a decrease in Notch activity, possibly via competing with other subunits, including NCT and PEN2, in the assembly of γ-secretase complexes and therefore its levels at the plasma membrane (Kaether et al., 2007; Spasic et al., 2007). To our knowledge, our observations provide the first evidence that Myc, which is a well-known target of Notch signaling (Klinakis et al., 2006; Satoh et al., 2004; Weng et al., 2006), could also modulate Notch activity by regulating Rer1 levels. Interestingly, past studies have shown that loss of Notch activity in clones of cells in the mammalian epithelium created super-competitor cells, which outcompeted the neighboring normal cells (Alcolea et al., 2014; Colom et al., 2020; Martincorena et al., 2018; Yokoyama et al., 2019). However, in contrast to this, Notch activity is required for the proliferation of cells in the *Drosophila* wing epithelium (Giraldez and Cohen, 2003; Herranz et al., 2008). Therefore, whether Notch activity in the *Drosophila* epithelium supports winner or loser cells remains unclear and whether lowered Notch activity in Myc-overexpressing cells is required for their super-competitive behavior remains to be investigated.

## MATERIALS AND METHODS

### Fly husbandry and fly strain

Standard food composition containing a cornmeal-sucrose-yeast was used to prepare the food. All crosses were maintained at 25L room temperature unless specifically mentioned. Egg collection, heat shock, dissection, immunostaining, and imaging were kept identical between the control and experimental setup. The following *Drosophila* stocks were used: *neoFRT 82B, rer1*^*KO*^ *in 3rd chr. (this paper), GFP-Rer1 on 2nd chr. and neoFRT82B, rer1*^*KO*^ *in 3rd chr. (this paper), neoFRT 82B, Ubi-mRFP in 3rd chr*., *(BDSC, 30555), FRT82B, UBI-GFP*.*nls, RpS3[Plac92] in 3rd chr*., *(BDSC, 5627), hs-FLP and UAS-GFP on 1st chr. and tubP-Gal4, neoFRT82B, tubP-Gal80 in 3rd chr*., *(BDSC, 86311), hh-Gal4 in 3rd chr. (Tanimoto et al*., *2000), UAS-Trip-rer1-RNAi on 2nd chr. (BDSC, 57435), UAS-GADD34 on 2nd chr. (BDSC, 576250), UAS-s6k*.*TE on 2nd chr. (BDSC, 6912)*.

### Generation of *rer1* mutants and genomic rescue fly-lines

To generate the *rer1* knock-out line, the fly line containing a P-element insertion upstream of the *rer1* coding sequence (y; ry506 P{SUPor-P}CG11857KG08816/TM3, Sb Ser) was obtained from the Bloomington *Drosophila* Stock Center Indiana (BDSC, 15137). The P-element imprecise excision was performed and the excision lines were screened by Polymerase chain reaction (PCR), using primers positioned in the terminal coding sequences of the neighboring genes. The Rer1 knock-out line, w;; FRT(neo)82B, ry506, *rer1*^*KO*^/TM6C,Sb,Tb was identified, with 1560bp deletion that covered the whole coding sequence of the *rer1* gene. To generate the GFP-rer1 (CG11857) BAC rescue construct, GFP was recombined to CG11857BAC genomic clone, CH322-101C14 (BACPAC resources) following the P[acman] method (Venken et al., 2009). The GFP-CG11857BAC was then inserted into VK18 (2L, 53B2) for genomic rescue.

### Antibodies

Larval wing imaginal discs were stained using the following antibodies: Rabbit anti-cleaved *Drosophila* Dcp-1 (1:300, Cell Signaling Technology), Rabbit-anti-p-eIF2α (1:300, Cell Signaling Technology). Mouse-anti-Cut (DSHB), Mouse-anti-NICD (DSHB), Mouse-anti-Beta gal (40-1A, DSHB). All these DSHB antibodies used are diluted into a 1:50 ratio. Fluorescent secondary antibodies used were Alexa-405, Alexa-488, Alexa-594, and Alexa-647 (Invitrogen) at 1:500 dilutions.

### Immunostaining

For antibody labeling, wandering third instar larvae were dissected and head complexes with wing imaginal discs were processed by following standard immunohistochemical procedures. Acridine orange staining was performed by dissecting the third instar larval wing imaginal discs in PBS (1X) followed by 2 minutes of incubation in 0.6 mg/ml acridine orange + PBS (1X) solution (Robbins and Marcus, 1963). Afterward wing discs were rinsed very briefly in PBS (1X) before mounting. For the detection of nascent protein synthesis, Click-iT™ Plus OPP Alexa Fluor™ 647 Protein Synthesis Assay Kit (Thermo Fisher Scientific) was used and the procedure was followed according to the manufacturer’s manual. Wing discs were mounted in Vectashield (Vector laboratories) on glass slides (1mm BlueStar micro slides). Staining and microscopy conditions for samples used respectively for comparisons were identical. Wing discs are oriented with the dorsal up and anterior left.

### Image acquisition and processing

Images of fixed samples were acquired using the 40x oil objective on a confocal microscope by Olympus (FV3000) with each slice (z-stack) equivalent to 1μm at both 1 and 5 times (X) optical zoom. Wing disc images were processed using ImageJ (ImageJ version 1.51j8, NIH) and Photoshop (Adobe Photoshop CS6 extended version 13.0 x64). Figures were made using Illustrator (Adobe Illustrator CS6 Tryout version 16.0.0).

### Statistical analysis

Wing disc pouch regions were taken for all the analysis. Clone size was measured as ‘‘total clone area per disc pouch area (%)’’ by analyzing all the clones in the pouch area of each genotype using ImageJ (ImageJ version 1.51j8, NIH) software. To calculate the fold change in p-eIF2α, OPP, GFP-Rer1, NICD, and Cut signals, the average pixel intensities of area of interest was divided by that of the corresponding wildtype area. All raw data was analyzed using Excel (Microsoft) and graphs were plotted using GraphPad (GraphPad Prism 8 version 8.0.2). Each raw data set is shown as a dot plot and the horizontal line represents the median. All ‘N’ numbers express biological replicates. Statistical analysis was performed by Welch’s t-test for single comparisons and by Ordinary one-way analysis of variance (ANOVA) with Tukey’s test for multiple comparisons. The significance level was set to p < 0.05. No statistical methods were used to predetermine the sample size. For the quantification of survival and development, 1st instar larvae were collected, and the numbers of survival larvae in different developmental stages were counted each day. All experiments were independently performed at least three times and were not randomized or blinded.

### Generation of clones

For genetic mosaic analysis, the FLP (Flippase)/FRT (Flippase recognition target) system (Theodosiou and Xu, 1998) was used to generate mosaic clones in the wing imaginal disc. Heat shock inducible FLP was expressed for the mitotic recombination in both mitotic clones and MARCM clones. Heat shock was given in a 37 °C water bath for 60 minutes at 48 hrs AEL (After egg laying). Larvae were then shifted to 25 °C and were dissected at 96 hrs AHS (After heat-shock) of the third-instar wandering stage.

### Drosophila genotypes

The following genotypes were used in this study:

Figure 1 A-A’ : *hs-*FLP*/+;* ; *FRT82B, Ubi-RFP/ FRT82B, Ubi-GFP*

Figure 1 B-C’ : *hs-*FLP*/+;* ; *FRT82B, Ubi-RFP/ neoFRT 82B, ry*^*506*^, *rer1*^*KO*^

Figure 1 D-D’ : *hs-*FLP*/+; GFP-rer1/+; FRT82B, Ubi-RFP/ neoFRT 82B, ry*^*506*^, *rer1*^*KO*^

Figure 1 F-I : *hs-*FLP*+;* ; *FRT82B, Ubi-RFP/ neoFRT 82B, ry*^*506*^, *rer1*^*KO*^

Figure 1 K-L’ : *hs-*FLP*/+;* ; *FRT82B, Ubi-GFP*.*nls, RpS3[Plac92]/ neoFRT 82B, ry*^*506*^, *rer1*^*KO*^

Figure 2 A-A’’,G-G’ : *hs-*FLP, *UAS-GFP/+;* ; *tubP-Gal4, neoFRT82B, tubP-Gal80/ neoFRT 82B, Ubi-mRFP*.*nls*

Figure 2 B-B’’,H-H’ : *hs-*FLP, *UAS-GFP/+;* ; *tubP-Gal4, neoFRT82B, tubP-Gal80/neoFRT 82B, ry*^*506*^, *rer1*^*KO*^

Figure 2 C-C’’,I-I’ : *hs-FLP, UAS-GFP/+; UAS-GADD34/+* ; *tubP-Gal4, neoFRT82B, tubP-Gal80/ neoFRT 82B, ry*^*506*^, *rer1*^*KO*^

Figure 2 D-D’’,J-J’ : *hs-FLP, UAS-GFP/+; UAS-s6k*.*TE/+* ; *tubP-Gal4, neoFRT82B, tubP-Gal80/ neoFRT 82B, ry*^*506*^, *rer1*^*KO*^

Figure 3 A : *hs-FLP, UAS-GFP/+;* ; *tubP-Gal4, neoFRT82B, tubP-Gal80/ neoFRT 82B, Ubi-mRFP*.*nls*

Figure 3 B : *hs-FLP, UAS-GFP/+; UAS-dmyc/+; tubP-Gal4, neoFRT82B, tubP-Gal80/ neoFRT 82B, Ubi-mRFP*.*nls*

Figure 3 C : *hs-FLP, UAS-GFP/+; UAS-dmyc/+; tubP-Gal4, neoFRT82B, tubP-Gal80/ neoFRT 82B, ry*^*506*^, *rer1*^*KO*^

Figure 3 D : *hs-FLP, UAS-GFP/+;* ; *tubP-Gal4, neoFRT82B, tubP-Gal80/ neoFRT 82B, ry*^*506*^, *rer1*^*KO*^

Figure 3 F-I’ : ; *GFP-rer1/UAS-dmyc; hh-Gal4, UAS-ER-RFP/+*

Figure 3 K-N : ; *GFP-rer1/+; hh-Gal4, UAS-ER-RFP/+*

Figure 3O-O’’ : *hs-*FLP, *UAS-GFP/+;* ; *tubP-Gal4, neoFRT82B, tubP-Gal80/ neoFRT 82B, Ubi-mRFP*.*nls*

Figure 3P-P’’ : *hs-*FLP, *UAS-GFP/+; UAS-dmyc/+; tubP-Gal4, neoFRT82B, tubP-Gal80/ neoFRT 82B, Ubi-mRFP*.*nls*

Figure 3Q-Q’’ : *hs-*FLP, *UAS-GFP/+; UAS-dmyc/+; tubP-Gal4, neoFRT82B, tubP-Gal80/ neoFRT 82B, ry*^*506*^, *rer1*^*KO*^

Figure 3R-R’’ : *hs-*FLP, *UAS-GFP/+;* ; *tubP-Gal4, neoFRT82B, tubP-Gal80/ neoFRT 82B, ry*^*506*^, *rer1*^*KO*^

Figure 4A-A’,F-F’ : *hs-*FLP, *UAS-GFP/+;* ; *tubP-Gal4, neoFRT82B, tubP-Gal80/ neoFRT 82B, Ubi-mRFP*.*nls*

Figure 4B-B’,G-G’ : *hs-*FLP, *UAS-GFP/+; UAS-dmyc/+; tubP-Gal4, neoFRT82B, tubP-Gal80/ neoFRT 82B, Ubi-mRFP*.*nls*

Figure 4C-C’,H-H’ : *hs-*FLP, *UAS-GFP/+; UAS-dmyc/+; tubP-Gal4, neoFRT82B, tubP-Gal80/ neoFRT 82B, ry*^*506*^, *rer1*^*KO*^

Figure 4D-D’,I-I’ : *hs-*FLP, *UAS-GFP/+;* ; *tubP-Gal4, neoFRT82B, tubP-Gal80/ neoFRT 82B, ry*^*506*^, *rer1*^*KO*^

## Author contributions

P.K.P and V.C. conceptualization; P.K.P., R.F.S., W.A. and V.C. investigation; P.K.P., W.A. and

V.C. resources and funding acquisition; P.K.P., W.A. and V.C. supervision; P.K.P. and V.C. writing original draft; R.F. S. and W.A. writing, review and editing.

## Acknowledgements

We thank IISER Bhopal for the fly facility and the DST-FIST facility for the confocal microscopy. The authors thank Lijun Jia for generating the rer1 mutant flies. We thank V.C. group members for their discussions on the project and manuscript.

## Funding

This work was supported by the Science and Engineering Research Board (SERB), Department of Science & Technology, Government of India (grant number: CRG/2021/004686). The laboratory of V. C. is also supported by intramural funds from IISER Bhopal and the Department of Biotechnology-EMR (grant number: BT/PR34467/BRB/10/1831/2019). P.K.P. was supported by a CSIR fellowship (09/1020/(0127)/2017-EMR-I).

W.A. acknowledges the financial support of the Vlaams Instituut voor Biotechnologie (VIB), KU Leuven (grant number: C14/21/095 and KA.20/085), the Fonds Wetenschappelijk Onderzoek (FWO) (grant number: I001322N), and the Stichting Alzheimer Onderzoek Belgie_ (grant number: #2020/0030)

## Conflict of interest

The authors declare no conflicts of interest.

## Notes

### Competing Interest Statement

The authors have declared no competing interest.

