## Supplemental Data for "Loss of Rer1 causes proteotoxic stress that drives cell competition and inhibits Myc-driven overgrowth"

#### *Drosophila genotypes*

Supplemental Figure 1E: *GFP-Rer1/UAS-rer1-RNAi; hh-Gal4/+*

Supplemental Figure 1F-F': *hh-Gal4/+*

Supplemental Figure 1G-G': *UAS-rer1-RNAi/+; hh-Gal4/+*

Supplemental Figure 1H-H': *hh-Gal4/+*

Supplemental Figure 1I-I': *UAS-rer1-RNAi/+; hh-Gal4/+*

Supplemental Figure 2A-B: *hs-FLP, UAS-GFP/+; ; tubP-Gal4, neoFRT82B, tubP-Gal80/ neoFRT 82B,* $ry^{506}$, $rer1^{KO}$

Supplemental Figure 2D: *hs-FLP, UAS-GFP/+; ; tubP-Gal4, neoFRT82B, tubP-Gal80/ neoFRT 82B, Ubi-mRFP.nls*

Supplemental Figure 2E: *hs-FLP, UAS-GFP/+; ; tubP-Gal4, neoFRT82B, tubP-Gal80/ neoFRT 82B,* $ry^{506}$, $rer1^{KO}$

Supplemental Figure 2F: *hs-FLP, UAS-GFP/ UAS-*$bsk^{DN}$*; ; tubP-Gal4, neoFRT82B, tubP-Gal80/ neoFRT 82B,* $ry^{506}$, $rer1^{KO}$

Supplemental Figure 3A-A''': *; GFP-rer1/+; ;*

Supplemental Figure 3B-B''': *; GFP-rer1/+; ;*

Supplemental Figure 4A-B: *hs-FLP/+; ; FRT82B, Ubi-RFP/ neoFRT 82B,* $ry^{506}$, $rer1^{KO}$

Supplemental Figure 4D-E: *hs-FLP/+; GFP-rer1/+; FRT82B, Ubi-RFP/ neoFRT 82B,* $ry^{506}$, $rer1^{KO}$

Supplemental Figure 5A-D: *hs-FLP, UAS-GFP/+; ; tubP-Gal4, neoFRT82B, tubP-Gal80/ neoFRT 82B,* $ry^{506}$, $rer1^{KO}$

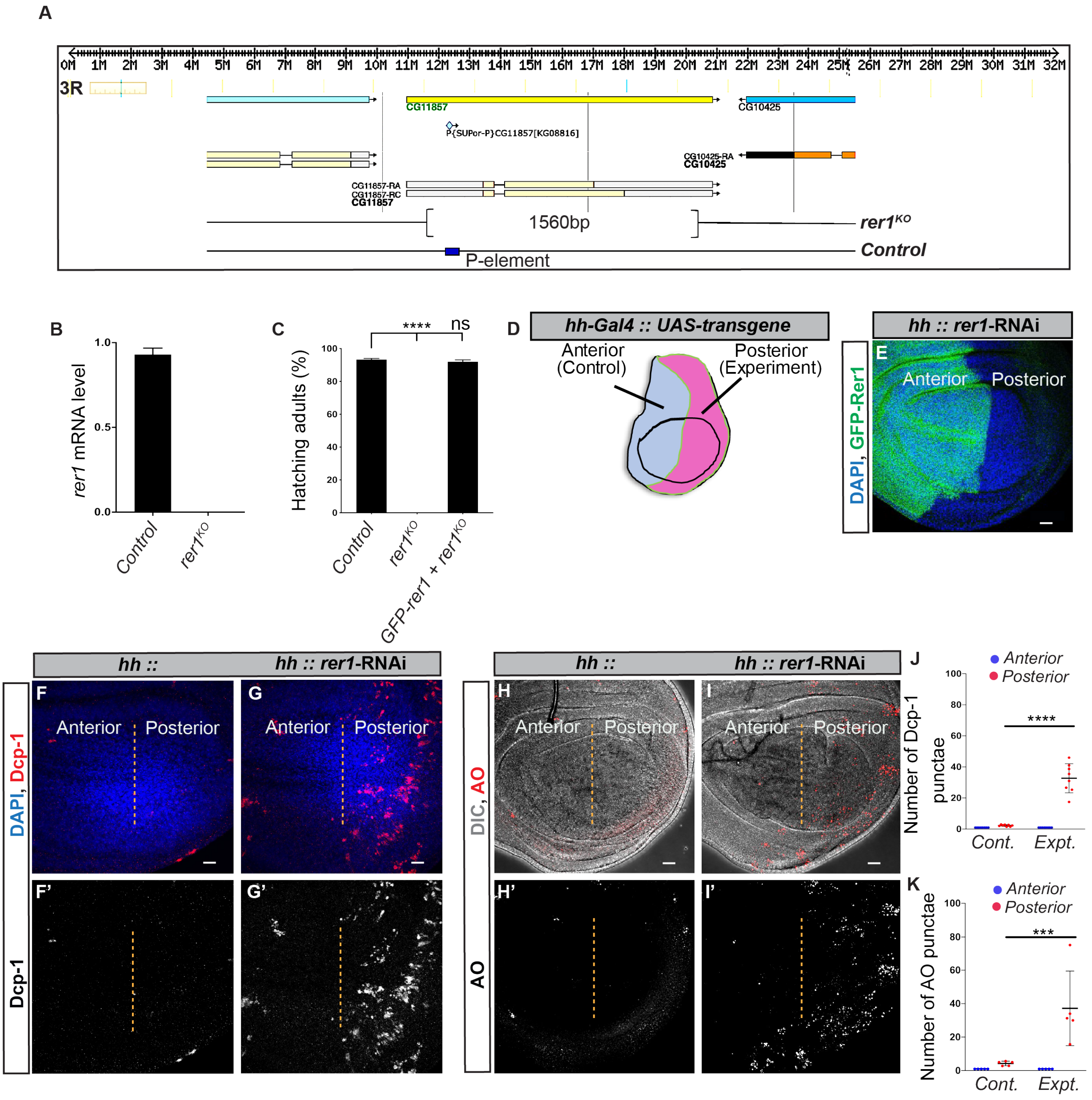


### *<u>Supplemental Figure 1</u>: Generation and characterization of rer1$^{KO}$ (rer1$^{-/-}$) fly line.*

*(A) Schematic representation of the rer1$^{KO}$ line. Upon imprecise excision of a p-element inserted in the coding sequence, a 1560bp deletion in the rer1 gene was obtained. The control line used contains the precise excision of the P-element retaining the whole coding sequence of the rer1 gene. (B) rer1 mRNA expression levels were measured by quantitative PCR. Bars show mean±SEM (N= 3 independent experiments). (C) Homozygous rer1$^{KO}$ (rer1$^{-/-}$) flies failed to hatch out. Most of them died before the pupae stage. Re-introducing the rer1 genomic fragment (GFP-rer1) rescued the phenotype, underscoring that they are caused by rer1 deficiency. Statistical analyses in C were performed using the Ordinary one-way ANOVA with*

*Dunnett's multiple comparison test. (D) Scheme of a wing disc illustrating anterior and posterior compartments in which transgene was expressed with a posterior specific Gal4. (E) GFP-Rer1 levels were analyzed in the posterior compartment upon hh-Gal4 mediated rer1 knockdown [GFP-rer1 + hh::UAS-rer1-RNAi] and compared to the anterior compartment. (F–G) Caspase activity was analysed in the posterior compartment upon hh-Gal4 mediated rer1 knockdown (G,G'), and compared to the posterior compartment of the control wing imaginal disc (F,F'). (H–I) Acridine Orange (AO) levels were analysed upon hh-Gal4 mediated rer1 knockdown (I,I')] and the posterior compartment was compared to the anterior compartment relative to the control wing imaginal disc (H,H'). (J-K) Quantification of Dcp-1 punctae (J) and AO punctae (K) in the wing pouch where the anterior compartment is the control, and the posterior compartment is the knockdown region.*

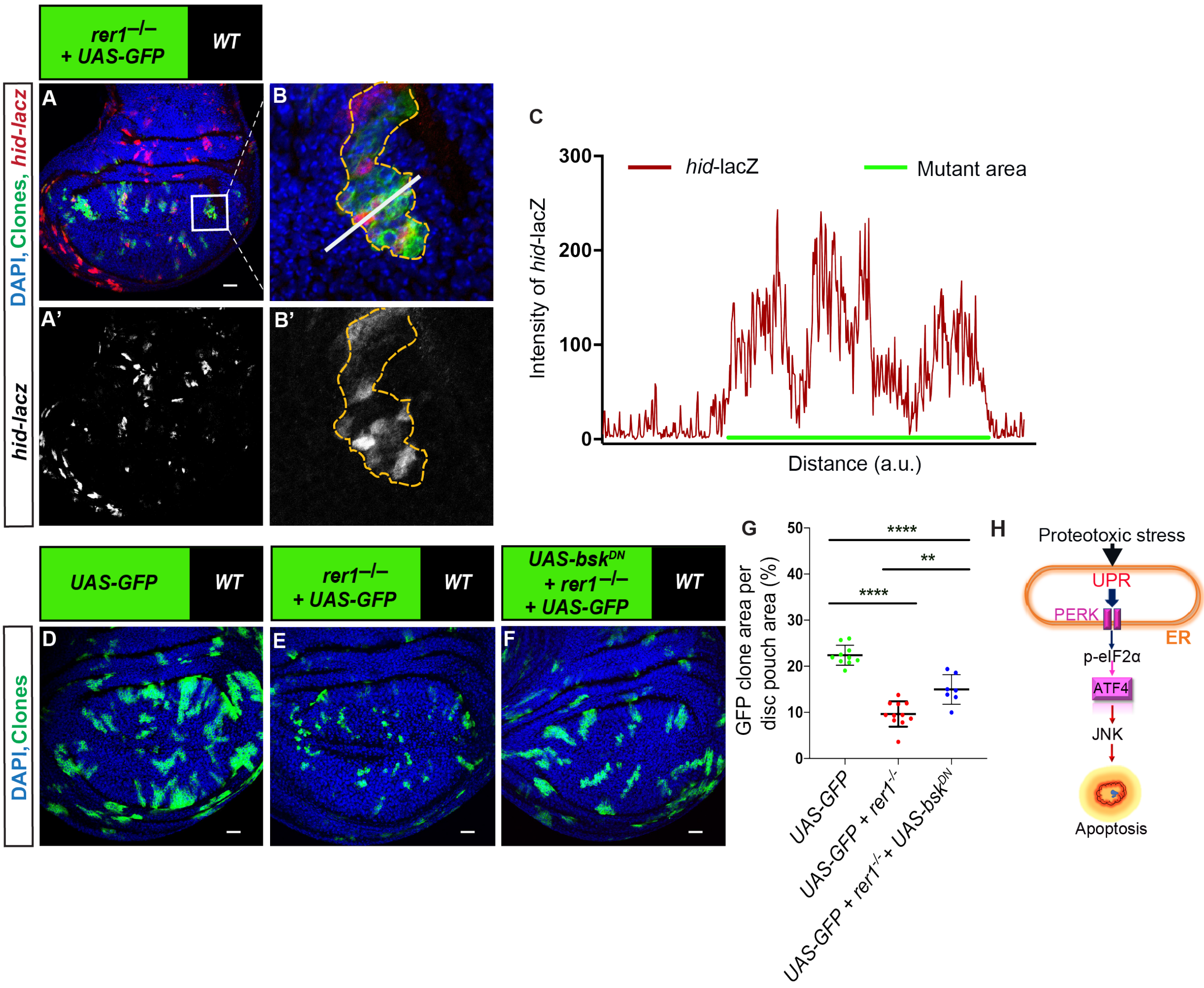


**<u>Supplemental Figure 2</u>: Involvement of JNK pathway in the elimination of rer1$^{-/-}$ tissues**

*(A) Third-instar discs containing hs-FLP-induced MARCM clones of UAS-GFP, rer1$^{-/-}$ genotypes, immuno-stained for the beta-galactosidase (red) to mark the hid-lacZ promoter activity; nuclei labeled in blue. (B) A magnified image of the inset (white box) in A. (C) Quantification of the intensity levels of hid-lacZ inside the GFP positive clones with respect to the nearby GFP negative control tissue along the line ROI region (white line) in B. (D-F) hs-FLP-induced MARCM clones of (D) UAS-GFP (rer1$^{+/+}$), (E) UAS-GFP, rer1$^{-/-}$ and, (F) UAS-GFP, rer1$^{-/-}$ + UAS-bsk$^{DN}$ genotypes with the nuclei labeled in blue in the third-instar wing discs epithelium. (G) Quantification of the relative size of GFP-labeled clone areas in UAS-GFP (rer1$^{+/+}$) (D, N=11), UAS-GFP, rer1$^{-/-}$ (E, N=11) and, UAS-GFP, rer1$^{-/-}$ + UAS-bsk$^{DN}$ (F, N=7). Statistical analyses in G were performed using the Ordinary one-way ANOVA with Tukey's multiple comparison test. (H) Diagram showing the involvement of PERK-peIF2α-JNK axis in resulting apoptosis upon proteotoxic stress. SB = 20 µm.*

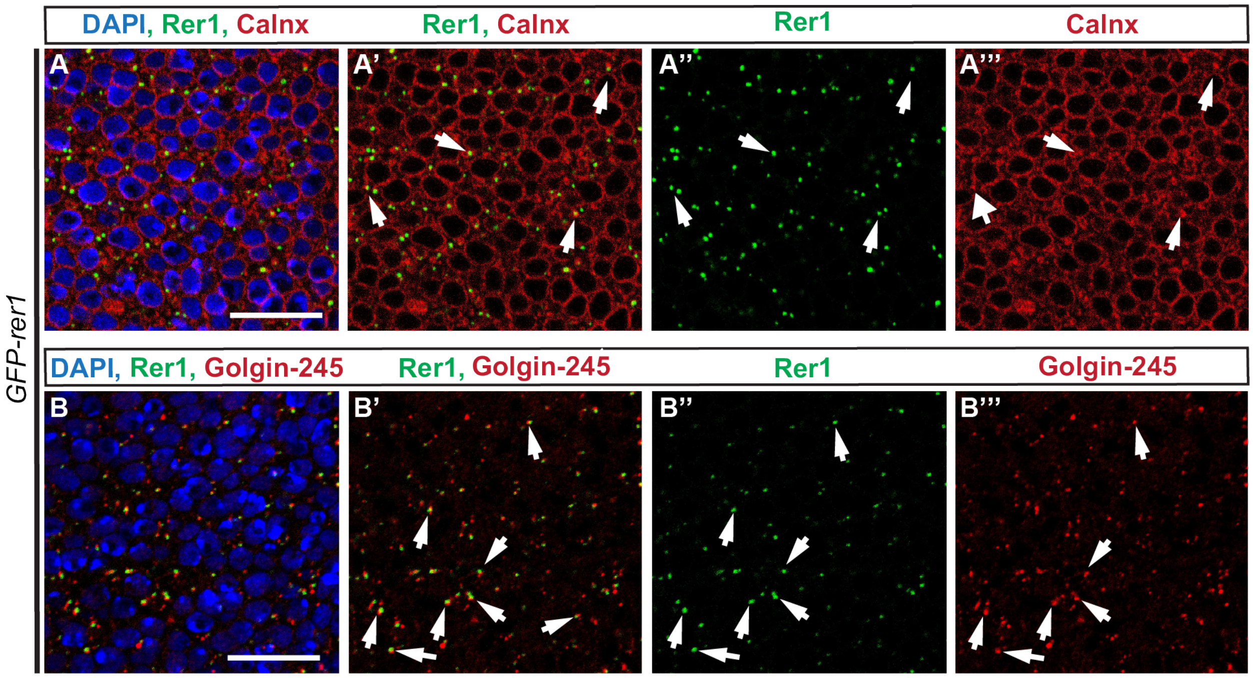


***<u>Supplemental Figure 3</u>: Colocalization study of Rer1 with ER and Golgi***

*(A-B) Colocalization study performed between Rer1 and ER in (A-A'''), staining with Calnexin; Rer1 and Golgi in (B-B'''), staining with Golgin-245. GFP tagged Rer1 protein (GFP-Rer1 transgene) was used. White arrows showed the colocalized punctae. SB = 20 um.*

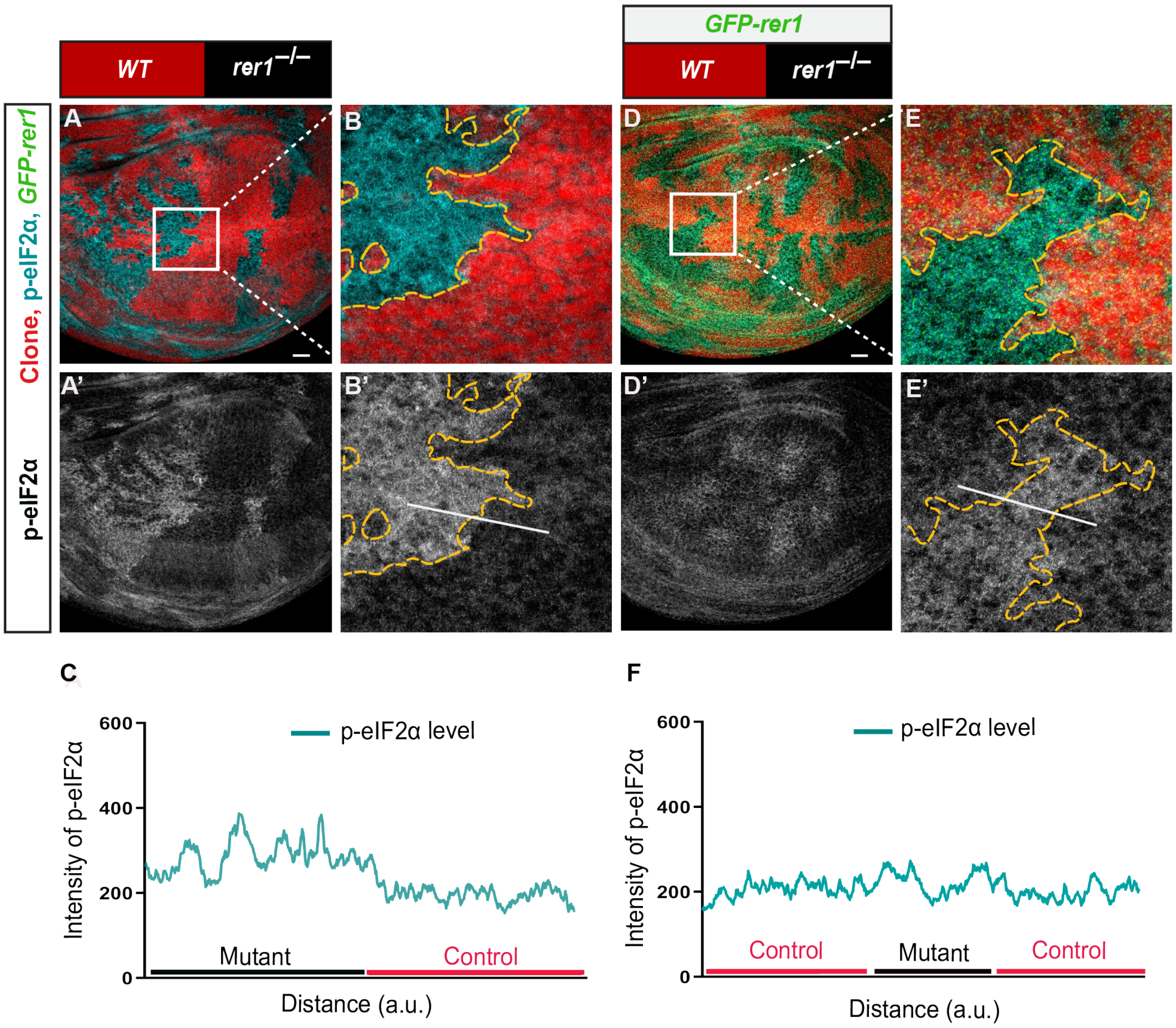


***<u>Supplemental Figure 4</u>: Ectopic expression of Rer1 rescues proteotoxic stress***

*(A) hs-FLP-induced mitotic clones of rer1$^{-/-}$ mutant tissues in third-instar larval wing epithelium, immuno-stained for the anti-p-eIF2α. (B) A magnified image of the inset (white box) in A. (C) Graph showing the intensity profile of p-eIF2α along the line ROI (white) in B' (N=5). (D) rer1$^{-/-}$ mutant clones in GFP-rer1 background stained with anti-p-eIF2α. (E) A magnified image of the inset (white box) in D. (F) Graph showing the intensity profile of p-eIF2α along the line ROI (white) in E' (N=9). Scale bars = 20 μm.*

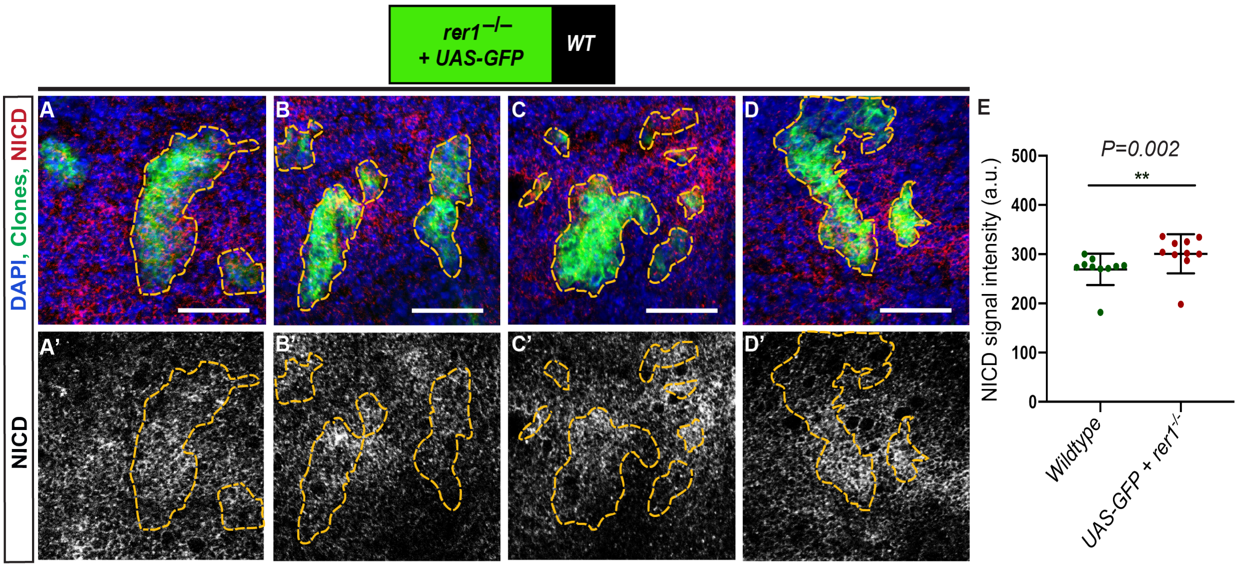


**<u>Supplemental Figure 5</u>: *Change of NICD levels in absence of Rer1***

*(A-D) hs-FLP mediated MARCM clones of UAS-GFP, rer1$^{-/-}$ genotypes showing the NICD expression levels in the third-instar wing disc epithelia. (E) Quantification of NICD signal intensity inside the GFP positive tissue compared to the GFP negative tissue (N=10; two-sided Wilcoxon signed-rank paired test) is shown. SB= 20 um.*
